# A ticket to ride: genetic structure and *kdr* mutations in *Aedes aegypti* populations along a road crossing the Amazon Rainforest in Amapá State, Brazil

**DOI:** 10.1101/2023.06.05.543667

**Authors:** Barbara S. Souza, Leticia F. Lima, Allan K. R. Galardo, Vincent Corbel, Jose Bento P. Lima, Ademir J. Martins

## Abstract

Insecticide resistance in *Aedes aegypti* is a major threat to the control of this mosquito species that transmits viruses that cause diseases like dengue, Zika, and chikungunya. One type of resistance is caused by alterations in the *Na_v_* gene, known as *kdr* mutations. In Brazil, different *kdr* haplotypes are present in *Ae. aegypti* and they may impede vector control operations based on pyrethroids. Although natural populations tend to accumulate genetic differences among isolated localities, this mosquito can actively and passively disperse, hitchhiked with human transportation. In this study, we investigated the genetic structure and *kdr* dispersion in *Ae. aegypti* populations in six localities of the Amapá State, Brazil located along a north-south transect of the Amazonian Forest. Genetic structure was assessed using 12 microsatellite loci in a fragment analysis sequencing procedure, and qPCR methods were used to detect the presence and frequency of three well known *kdr* mutations (V410L, V1016I and F1534C). We found a high prevalence of *kdr* alleles in all localities, indicating that *kdr* is spreading in the Amapá State. The microsatellite analyses suggested a certain level of differentiation among the mosquito populations, dividing them into two well-defined clusters, as evidenced by Bayesian and DAPC analyses. The population from Oiapoque (located in the north along the border with French Guiana) had the highest *kdr* frequencies and the highest genetic differentiation compared to the other localities. Our findings suggest that there is genetic structure among *Ae. aegypti* from the Amapá State, but with some level of passive gene flow between population clusters. The study highlights the importance of continued surveillance of *Ae. aegypti* populations to monitor the spread of insecticide resistance and inform on vector control strategies.

**Author summary:** Amapá, a State in northern Brazil, is crucial as a gateway for diseases due to its border with French Guiana. One notable example was chikungunya, transmitted by the *Aedes aegypti* mosquito, also responsible for dengue, Zika, and yellow fever viruses. Chemical insecticides have been the primary means of control, however, resistance to these compounds is spreading among *Ae. aegypti* populations, compromising their effectiveness. Mutations known as “*kdr*” contribute to resistance to pyrethroid insecticides. Our study focused on these mutations and genetic differentiation in *Ae. aegypti* populations from six cities along the road nestled amidst the Amazonian Forest, linking the capital, Macapá, to Oiapoque on the Brazil-French Guiana border. The results revealed high frequencies of *kdr* mutations in all populations, indicating probable resistance to pyrethroids. Genetic analysis showed two distinct groups among the mosquitoes, with evidence of mosquito flow, particularly between Macapá and Oiapoque, facilitated by human transportation. Monitoring insecticide resistance and mosquito migration is essential for effective vector control strategies. Understanding these factors enables us to combat mosquito-borne diseases and safeguard public health.

## Introduction

Arboviruses (*Arthropod-borne virus*) infections, such as dengue, Zika, chikungunya and yellow fever are causing severe impacts on global health, especially in tropical low-income countries [1]. In Brazil, dengue is considered as hyperendemic, with regional and seasonal circulation of the four serotypes (I-IV), but other newly emerged viruses such as Zika (ZIKV) and chikungunya (CHIKV) are also circulating [2]. In 2022, Brazil reached the highest number of deaths caused by dengue fever (1,016 deaths) ever recorded in the country [3], and the recent increase in yellow fever cases in regions bordering urban centers [4] highlights the imminence of re-urbanization of this disease [5]. As there are no effective vaccines available for most of these arboviral diseases (excluding yellow fever), controlling the vector – the mosquito *Aedes aegypti* (Diptera, Culicidae: Linnaeus, 1762) – is the method of choice for reducing the risk of arbovirus transmission.

Chemical control is still the cornerstone of any vector borne-diseases control programme worldwide [6,7]. Historically, insecticides have played an essential role in the decline of various diseases, including malaria and dengue [8]. However, the massive and repeated use of chemicals for decades has favored the spread of insecticide-resistance in *Aedes* mosquitoes [9]. Currently, insecticide resistance (IR) is seen as one of the most important threats for the control of diseases caused by viruses transmitted by *Aedes,* as it may reduce the efficacy of chemical-based vector control interventions [10,11]. In Brazil, nationwide insecticide resistance monitoring (MRI) has reported strong resistance of *Ae. aegypti* to all pesticides used by the Ministry of Health, such as the larvicide temephos (organophosphate), the insect growth regulator pyriproxyfen, and the adulticides deltamethrin (pyrethroid) and malathion (organophosphate) [12,13].

Among the genetic alterations involved in IR, the most common are single nucleotide changes in the voltage-gated sodium channel (*Na_v_*), which cause resistance to the knockdown effect of pyrethroids, and are therefore called the *kdr* mutations (knockdown resistance) [14]. In Brazil, at least three *kdr* mutations have been reported in *Ae. aegypti*: a substitution of Valine to Isoleucine at position 1016 (V1016I), a Phenylalanine to Cysteine at position 1534 (F1534C), and a Valine to Leucine at position 410 (V410L) [15,16]. These mutations were classified as *Na_v_R1* and *Na_v_R2*, where *Na_v_R1* is a haplotype containing the F1534C *kdr* mutation only while *Na_v_R2* exhibits F1534C + V410L + V1016I mutations [17]. These alleles give pyrethroid resistance under a recessive trait, and the *Na_v_R2* causes the highest level of resistance to deltamethrin [18]. Other SNPs, such as V1016G, S989P and T1520I, are common in Asian populations of *Ae. aegypti* [19,20] but they are absent in Brazil and neighboring countries.

Monitoring the presence and spread of *kdr* mutations through molecular methods is relevant because pyrethroids are widely used for vector control and household domestic purposes [21,22,16]. Additionally, the analysis of the genetic structure and gene flow among vector populations can contribute to a better understanding of the evolutionary forces driving the dispersion of resistance alleles [23].

In Brazil, differences in the genetic structure and spatial distribution of pyrethroid (*kdr*) resistance have been reported in *Aedes aegypti* with at least three well defined clustered regions within the country [16]. The Amapá State (AP), located in the Northern part of the country, bordering the French Guiana, is an important gateway between Brazil and the Caribbean to arboviruses and vector populations with distinct characteristics, including insecticide resistance [24]. By the way, this was one of the entrance doors for the CHIKV virus in 2015 in Brazil [25]. In addition, the geographical aspects of the region raise relevant questions for population genetic studies since locations infested with *Ae. aegypti* are relatively isolated from each other’s due to the vast Amazonian vegetation. In AP, *Ae. aegypti* from Oiapoque (the city bordering the French Guiana) is among the Brazilian mosquito populations exhibiting the highest levels of resistance to deltamethrin, temephos and malathion, while the *Ae. aegypti* population from the AP State capital, Macapá city, is far less resistant [12,26,27]. These cities are 577 km distant, connected by a road (BR-156) that crosses the dense Amazon Rainforest. The frequency of *kdr* alleles is distinct between them, with a predominance of *kdr Na_v_R1* and *kdr Na_v_R2* in Macapá and Oiapoque, respectively [24]. The presence and frequency of these *kdr* alleles in other cities of the AP State is however unknown.

In this study, we conducted mosquito collection in cities located along the BR-156 road of the Amapá State to assess the genetic structure and the spatial distribution of *kdr* mutations in *Ae. aegypti* along a South-North transect. The study was carried out to better understand local adaptation of dengue vector populations in the Amapá State and guide national authority in the selection of the most judicious insecticides to use for vector control.

## Materials and methods

### *Ae. aegypti* collections

The field team of *Laboratório de Entomologia Médica do Instituto de Pesquisas Científicas e Tecnológicas do Estado do* Amapá (IEPA) installed 50 eggtraps in each of the six Amapá cities: Oiapoque (OIA) (03°49’53’’ N, 51°50’07’’ W), Calçoene (CAL) (02°29’53’’ N, 50°56’59” W), Tartarugalzinho (TTZ) (01°30’21” N, 50°54’41’’ W), Ferreira Gomes (FGO) (00°51’14” N, 51°11’39” W), Porto Grande (PGR) (00°42’16’’ N, 51°24’35’’ W) and Macapá (MAC) (00°02’04” N, 51°03’60” W) (Fig 1), following the methodology developed by the Ministry of Health [13]. Collections were made in August (OIA, CAL, TTZ, FGO, PGR) and October/2020 (MAC). The eggtraps in CAL, FGO, PGR and TTZ were randomly distributed in the whole city. In MAC and OIA, the eggtraps were preferentially placed on the sidelines of BR-156 road. The eggs were stimulated to hatch in IEPA laboratory, and the resulting adults (F0) were packed in silica gel and shipped to *Laboratório de Biologia, Controle e Vigilância de Vetores* (LBCVIV) at *Instituto Oswaldo Cruz* (IOC/Fiocruz) for further analysis.

**Fig 1.**
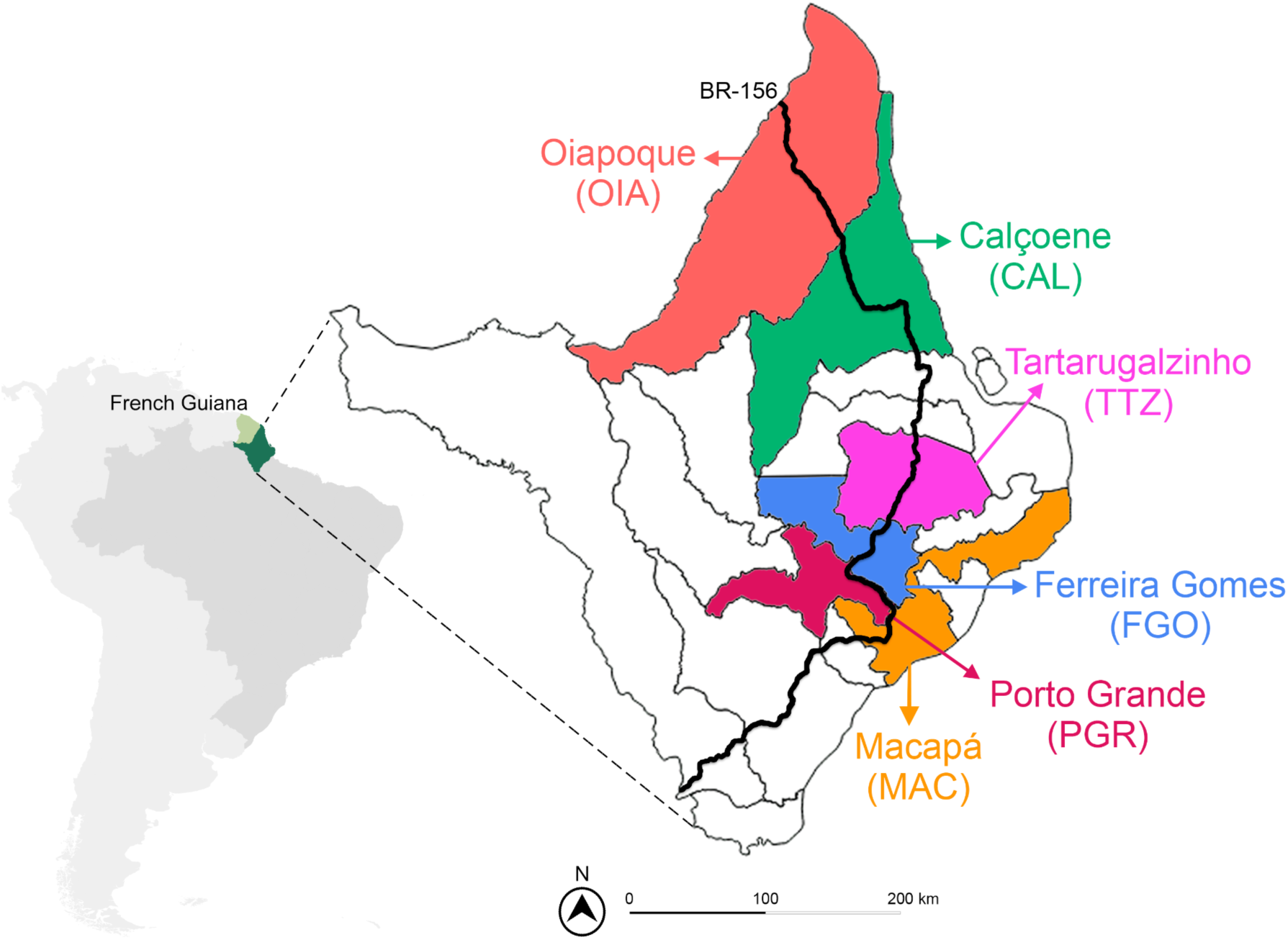
Map showing the location of Amapá State, Brazil, and French Guiana, France. The delimitations of Amapá cities are also shown including the cities where *Aedes aegypti* were collected (in colour). The BR-156 road is indicated by a black line.

### *Kdr* genotyping

We extracted DNA from male mosquitoes from the F0 generation (from egg collection), individually titrated with TNES buffer, as described elsewhere [28]. We use males to avoid amplifying the DNA of sperm from inseminated females’ spermathecae. The DNA of each sample was quantified in a Nanodrop One C spectrophotometer (Thermo Scientific) and aliquoted at 20 ng/µL in ultra-pure water. About 45 individuals from each population were genotyped for the *kdr* SNPs V410L, V1016I and F1543C through real-time TaqMan qPCR approach, essentially as described elsewhere [16] (See S1 Table for primers and probes sequences). The genotyping callings of each SNP were resolved by the online software Genotype Analysis Module v3.9 (Thermo Fisher), setting the endpoint to CT 40. Allelic and genotypic frequency charts were performed on GraphPad Prism v9.2.0 (www.graphpad.com). We considered the three SNPs (410+1016+1534) to determine the genotype of each sample [16,20].

### *Na_v_* Sequencing

We amplified and sequenced fragments corresponding to the IS6, IIS6, and IIIS6 *Na_v_* segments of elected samples to determine the respective *kdr* haplotypes circulating in *Ae. aegypti* populations from Amapá. We selected nine samples genotyped as homozygous to *kdr Na_v_R1* or *kdr Na_v_R2* alleles (see Results) from MAC and OIA. For the PCR amplification, we used the kit Phusion High-Fidelity PCR (New England, Biolabs), containing Phusion Buffer 1X, DMSO 3%, the respective primers pair (S2 Table) 0.5 µm each and ultra-pure water q.s. 25 µL. The thermal-cycle conditions were 98°C/30″ in the first step, followed by 35 cycles at 98°C/10″ for denaturation, 60°C (IS6), or 57°C (IIS6), or 61°C (IIS6)/15″ for primers annealing, and 72°C/30″ for the Polymerase amplification, followed by a final extension step at 72°C/7’. The amplicons were purified with the Qiaquick PCR Purification kit (Qiagen), according to the conditions provided by the manufacturer, and subjected to the sequencing reaction with the Bigdye Terminator V3.1 kit (Invitrogen, Thermo Fisher) using 1 µL of the purified amplicon and 1 µM of one of the respective primers. The sequencing reaction products of both strands of each sample were sent to the Fiocruz DNA Sequencing Facility (an ABI 3730 equipment). The sequences were analyzed using Geneious v9.1.8 [29].

### Microsatellites genotyping

The same DNA samples used for *kdr* genotyping were also genotyped for 12 well-characterized and polymorphic microsatellite loci that are widely used in studies of *Ae. aegypti* populations from the American continent: AC1, AC2, AC4, AC5, AG1, AG2, AG5, CT2, A1, A9, B2 and B3 (S3 Table) [30,31]. We performed the reactions with the Type-It PCR kit (Qiagen), according to a protocol standardized by Brown et al. 2011 [30], using multiplex primers, originally proposed by Schuelke 2000 [32], using an M13 tail at the primers 5’ end and marked with FAM or HEX fluorescence (S3 Table). For each reaction, we used the Type-It Multiplex PCR Master Mix (Qiagen) 1x, each forward primer at 0.025 μM, each reverse primer at 0.25 μM, each probe at 0.5 μM, 1 µL of DNA (20 ng) and ultra-pure water q.s. 10 µl. The thermocycling conditions were 94°C/10’ followed by 35 cycles of 94°C/30”, 54°C/30” and 72°C/30”, followed by the final step of 72°C/5’. The product of each PCR was diluted 1:10, and 1 µL of the product was used for genotyping. Each amplicon received 0.5 μM the dye size standard GeneScan 500 LIZ (Applied Biosystems) and was submitted to the Genotyping/ Fragment Analysis Facility at Fiocruz (equipment 3130xl Genetic Analyser, Applied Biosystems).

### Data analysis

We used the software *Geneious* 9.1.8 [29] with the *plug-in* developed for ABI fragment analyses (*Geneious Microsatellite Plugin*) to obtain the genotype callings of all microsatellite loci, which were exported in a .csv file. We tested the loci for Hardy-Weinberg equilibrium (HWE) and linkage disequilibrium in the Genepop v4.7.5 software, with significance levels adjusted by Bonferroni correction [33,34]. The presence of null alleles in each locus was verified with Microchecker Software v2.2.3 [35]. The genetic diversity parameters: average number of different alleles (Na), number of effective alleles (Ne), private allelic richness (Np), expected heterozygosity (He), observed heterozygosity (Ho), endogamy coefficients (Fis) and the number of migrants (Nm) for each population were estimated using the software Genetic Analysis in Excel (Genalex) v.6.503 [36]. This same software was used for analysis of molecular variance (AMOVA). The allelic richness (R) was calculated with the HP-Rare v1.1 software [37]. We estimated the magnitude of genetic differentiation with paired Fst values using Arlequin v3.5.2 [38] and Freena [39], respectively, both with 10.000 permutations. To test the assumption of isolation by distance (IBD), we run the Mantel test, with Arlequin v3.5.2 with 10.000 permutations [38], correlating genetic and geographic data (geographical distance per km x Fst).

We also used the Structure v2.3.4 software [40] to assess the population’s genetic structuring based on the number of genetic clusters (K). The best K value was obtained based on ten independent runs with 500,000 Monte Carlo chains (MCMC) iterations, excluding 20% of the initial chains (burn-in). The output data was analyzed in the Structure Harvester 2.3 software [41] to determine the best number of genetic clusters (K) based on an analysis of the ΔK chart [42]. Finally, we submitted the output files to Clumpp v1.1.2 [43] and Distruct V1.1 [44] software to plot the genetic structure of the evaluated populations. A multivariate statistical analysis (discriminant analysis of principal components -DAPC) was performed with the Adegenet package [45,46] in the R platform [47].

## Results

### *Kdr* genotyping

We genotyped 272 *Ae. aegypti* mosquitoes for the three *kdr* SNPs V410L, V1016I and F1534C. In total, we found eight genotypes (Fig 2): the six genotypes expected by the combination of wild-type *Na_v_S* (VVF), *kdr Na_v_R1* (VVC) and *kdr Na_v_R2* (LIC) alleles, and two additional. The genotype R2X1 (LL+VI+CC), composed of *kdr Na_v_R2* and the herein called *kdr Na_v_X1* (LVC) alleles, was found in OIA and CAL. We could not accurately determine the allelic composition of the genotype Y (LL+VI+FC), because it may be composed of LVF/LIC or LIF/LVC. This genotype was exclusively found in OIA. We named the possible alleles LFV and LIF as *kdr Na_v_X2* and *kdr Na_v_X3*, respectively (S4, S5 Tables and Fig 2).

**Fig 2.**
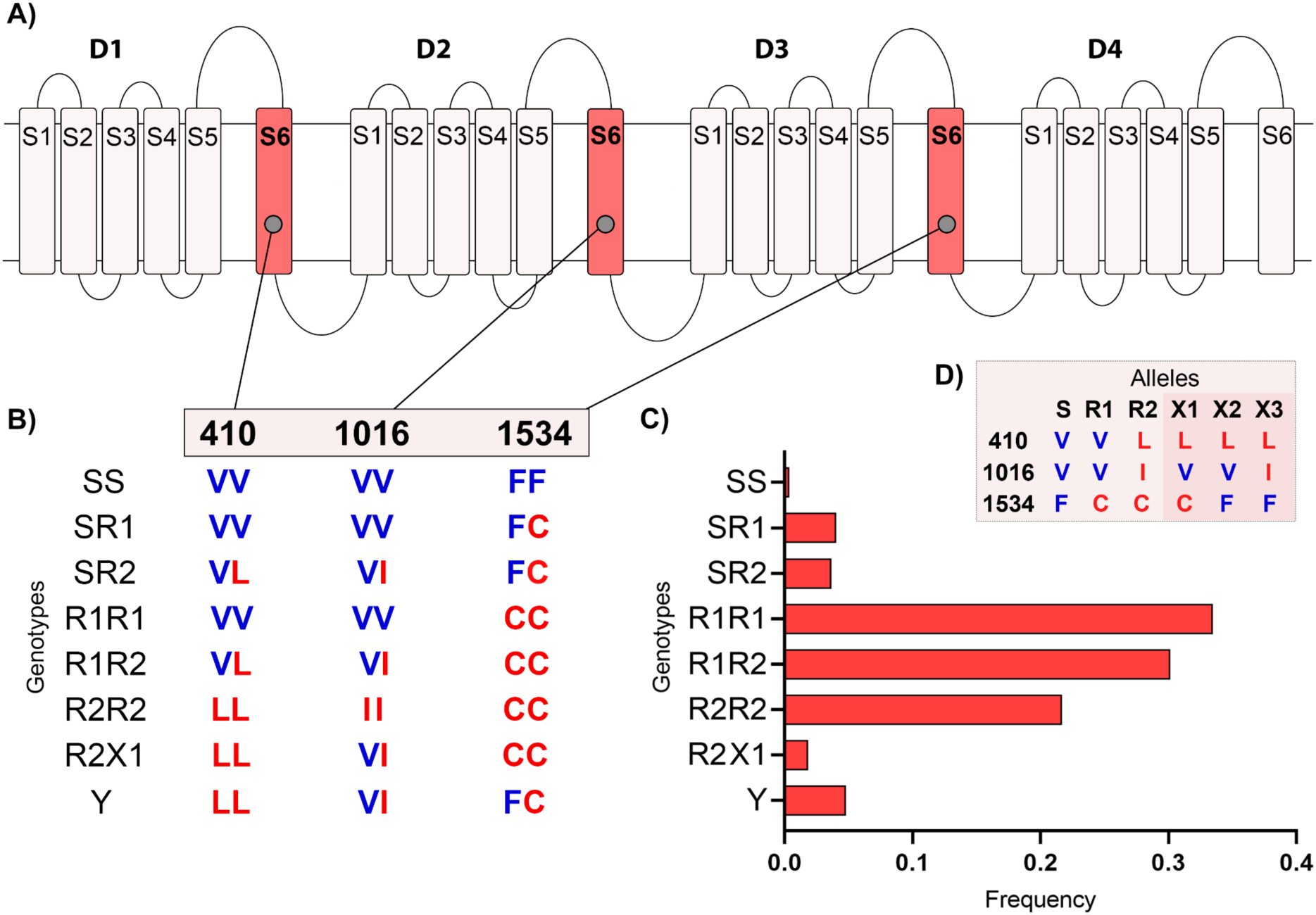
*Kdr* genotypes in *Aedes aegypti* from the Amapá State, Brazil. A) scheme of the voltage-gated sodium channel (*Na_v_*), indicating the four domains (D1-D4), each with six transmembrane segments (S1-S6). B) The genotypes observed in each of the three SNP sites in IS6, IIS6 and IIIS6 *Na_v_* segments. The wild-type and the *kdr* genotypes are in blue and red, respectively. C) The total *kdr* genotypic frequencies, considering all samples (272 mosquitos genotyped for V410L, V1016I and F1534C) from the Amapá State. D) Possible alleles that constitute the genotypes found here. The “Y” genotype is obtained by combining R2/X2 or X1/X3 alleles.

In general, the lowest genotype frequency found in the *Ae. aegypti* populations was the wild-type homozygous SS (VV+VV+FF), only observed in MAC (2.3%). The most frequent genotype was R1R1 (VV+VV+CC), ranging from 2.2% in OIA to 65.1% in MAC. All populations showed high frequencies of other “resistant genotypes” (i.e. R1R1, R1R2, and R2R2), whereas their sum ranged from 63% in OIA to 100% in PGR. MAC was the only population without the homozygous R2R2 genotype (R1R1 + R1R2 = 76.7%). The genotype R2X1 was observed in CAL (8.7%) and OIA (2.2%). The genotype Y (LL+VI+FC) was found only in OIA, although at a relatively low frequency (28.3%) (S4 Table and Fig 2). The populations showed opposite trends regarding to the *kdr* allelic frequencies according to their location along the South-North transect. For example, MAC presented the higher *kdr Na_v_R1* (77.9%) and the lower *kdr Na_v_R2* (9.3%) frequencies, while OIA showed the lower *kdr Na_v_R1* (15.2%) and the higher *kdr Na_v_R2* (52.2%) frequencies (S5 Table). The genotypic frequencies of each population are presented in S1 Fig.

### *Kdr* haplotype sequences

We amplified and sequenced the corresponding IS6, IIS6 and IIIS6 *Na_v_* fragments of some homozygous R1R1 (4 MAC and 1 OIA) and R2R2 (4 OIA) mosquitoes. The IS6, IIS6 and IIIS6 sequences of R1R1 samples were similar to previously published sequences (GenBank accession numbers: LC557528, MN602762, MN602780, respectively). The R2R2 sequences of samples from OIA, IS6, IIS6, and IIIS6 fragments were also similar to known sequences (GenBank accession numbers: KY747530, MN602754, MN602780, respectively).

### Microsatellite analyses

We genotyped 12 microsatellite loci in a total of 288 *Ae. aegypti* divided into six populations from the Amapá State. We observed a total of 60 alleles, varying from two (AC4 and B2) to 17 (Ag2) (S6 Table). Some markers were not under HWE, even after Bonferroni corrections, in some populations: AG2 (CAL, OAI), A9 (TTZ, CAL), A1 (FGO), AC1 (FGO, TTZ), AG5 (FGO, CAL) and AG1 (FGO) (S7 Table). In this case, the populations showed a Fis>0 (Table 1 and S7), indicating a heterozygous deficit. Although the LD test showed 33 tests significant out of the 396 evaluated combinations (8.3%), none loci pairs were consistently correlated in all populations after Bonferroni correction (S8 Table).

**Table 1.**
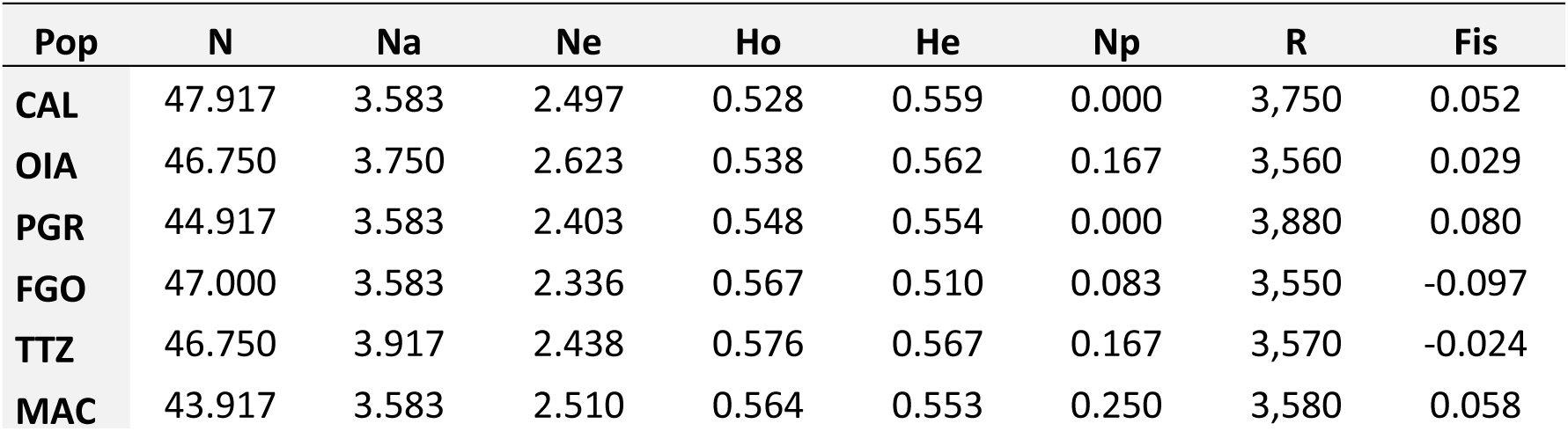
Genetic diversity of six *Aedes aegypti* populations from Amapá State, Brazil, based on the analysis of 12 microsatellites.

| Pop | N | Na | Ne | Ho | He | Np | R | Fis |
| --- | --- | --- | --- | --- | --- | --- | --- | --- |
| CAL | 47.917 | 3.583 | 2.497 | 0.528 | 0.559 | 0.000 | 3,750 | 0.052 |
| OIA | 46.750 | 3.750 | 2.623 | 0.538 | 0.562 | 0.167 | 3,560 | 0.029 |
| PGR | 44.917 | 3.583 | 2.403 | 0.548 | 0.554 | 0.000 | 3,880 | 0.080 |
| FGO | 47.000 | 3.583 | 2.336 | 0.567 | 0.510 | 0.083 | 3,550 | -0.097 |
| TTZ | 46.750 | 3.917 | 2.438 | 0.576 | 0.567 | 0.167 | 3,570 | -0.024 |
| MAC | 43.917 | 3.583 | 2.510 | 0.564 | 0.553 | 0.250 | 3,580 | 0.058 |

### Genetic diversity and differentiation

Table 1 shows the genetic diversity in the *Ae. aegypti* populations from the Amapá State. The average allelic diversity varied from 3.58 (MAC) to 3.91 (TTG), the average number of effective alleles (Ne) from 2.33 (FGO) to 2.62 (OIA), the number of private alleles per population (Np) from 0 (CAL and PGR) to 0.25 (MAC), and the number of allelic richness (R) from 3.56 (CAL) to 3.88 (TTZ). The genetic diversity of a mosquito population usually is positively related to the expected heterozygosity (He) and observed heterozygosity (Ho) values. Here the He varied from 0.510 to 0.567 and the Ho varied from 0.528 to 0.576 (Table 1).

Regarding the genetic differentiation as measured by the Fst, the values of the pairs ranged from 0.004 (PGR-TTZ) to 0.084 (OIA-PGR) (Table 2 and S9). Except for the pairs OIA-FGO (0.818) and OIA-PGR (0.884), the number of migrants (Nm) values were greater than one, indicating a certain degree of gene flow between the *Ae. aegypti* populations along the BR-156 road. The pairs associated with OIA showed the highest Fst and, conversely, the lowest Nm values, which can be justified by the fact that Oiapoque is the city located in the extreme north of the BR-156 road. Interestingly, OIA-MAC showed the highest Nm value among all pairs with OIA, although Macapá and Oiapoque are the most distant cities. The AMOVA showed Fis and Fit values of 0.92 and 1.57 (both p <0.001), respectively (S10 Table), indicating a moderate differentiation within the populations. At least 55% of the genetic distance among the populations may be attributed to isolation by distance (IBD), according to the Mantel test (0.549, p>0.031).

**Table 2.** Genetic differentiation (Fst) and number of migrant (Nm) indexes of pairwise comparison of *Aedes aegypti* from Amapá State, Brazil.

| Fst/Nm | CAL | OIA | PGR | FGO | TTZ | MAC |
| --- | --- | --- | --- | --- | --- | --- |
| CAL | - | 1,009 | 2,238 | 1,276 | 2,579 | 1,581 |
| OIA | 0,076 | - | 0,884 | 0,818 | 1,167 | 1,229 |
| PGR | 0,031 | **0,084** | - | 8,390 | 53,037 | 3,696 |
| FGO | 0,066 | 0,074 | 0,021 | - | 6,540 | 4,597 |
| TTZ | 0,034 | 0,067 | **0,004** | 0,016 | - | 3,569 |
| MAC | 0,040 | 0,050 | 0,026 | 0,031 | 0,019 | - |

\* Bellow diagonal: Fst values; above diagonal: Nm values. Marked in bold are the highest and lowest Fst values. CAL: Calçoene; OIA: Oiapoque; PGR: Porto Grande; FGO: Ferreira Gomes; TTZ: Tartarugalzinho; MAC: Macapá.

### Genetic structure

The Bayesian analysis conducted on the mosquito populations from Amapá suggests that *Ae. aegypti* is most likely divided into two genetic clusters (K=2) (S2 Fig). We designed the structure plot considering K=2 and K=3 (Fig 3A). In both scenarios, TTZ, FGO, PGR and MAC are the most homogeneous populations. With K=2, OIA and CAL seem more related; with K=3, OIA stands out as a more isolated population. The pie charts with the cluster frequencies in each population over the map (Fig 3B) facilitate observing the populations’ genetic structure along the BR-156 road. For example, with K=2, we observed that the most homogeneous populations were OIA and FGO, with 95.7% of the genetic diversity assigned back to cluster 2 in OIA while 95.5% to cluster 1 in FGO. In addition, the DAPC analysis plot evidenced threegroups, where each OIA and CAL formed isolated groups, and TTZ, FGO, PGR and MAC were all mixed in a third group (Fig 3C). These genetic structure analyses complemented the genetic differentiation indexes, suggesting that OIA and CAL are more structured than the other populations. This trend can be explained by the likely higher gene flow among the mosquitoes from TTZ, FGO and PGR and the capital MAC compared to OIA, which is a distant and isolated city located in the border area with French Guiana.

**Fig 3.**
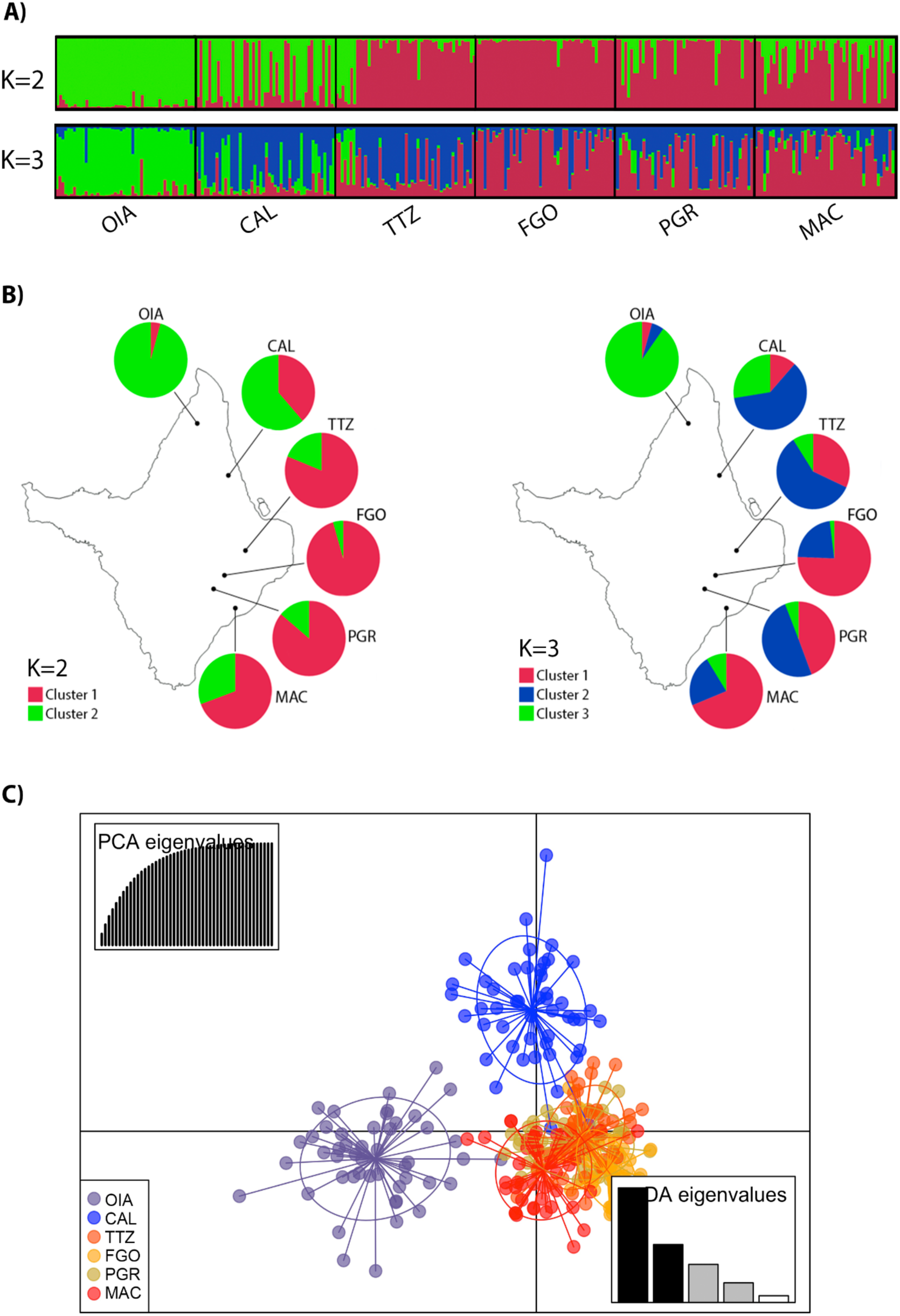
Genetic structure of *Aedes aegypti* populations from Amapá State, Brazil, based on 12 microsatellite markers. A) Structure plot: Bayesian clustering analysis where each bar represents the inferred ancestry of each individual with two (K=2) or three (K=3) genetic clusters. B) Pie charts of the global inferred ancestry value of each population with K=2 and K=3. C) Discriminating analyses of principal components (DAPC) plot, with the dots and colors representing individuals and groups, respectively. Eigenvalues indicate the number of principal components that best explain the differences between individuals (4 components were indicated for the 6 populations).

### *Kdr* and genetic structure

Considering the genetic structure based on the SSR analyses into two clusters (K=2), the populations from clusters 1 and 2 presented the *kdr* alleles *Na_v_R1* and *Na_v_R2*, whilst the additional *kdr* alleles were present only in the populations from cluster 2 (Fig 4).

**Fig 4.**
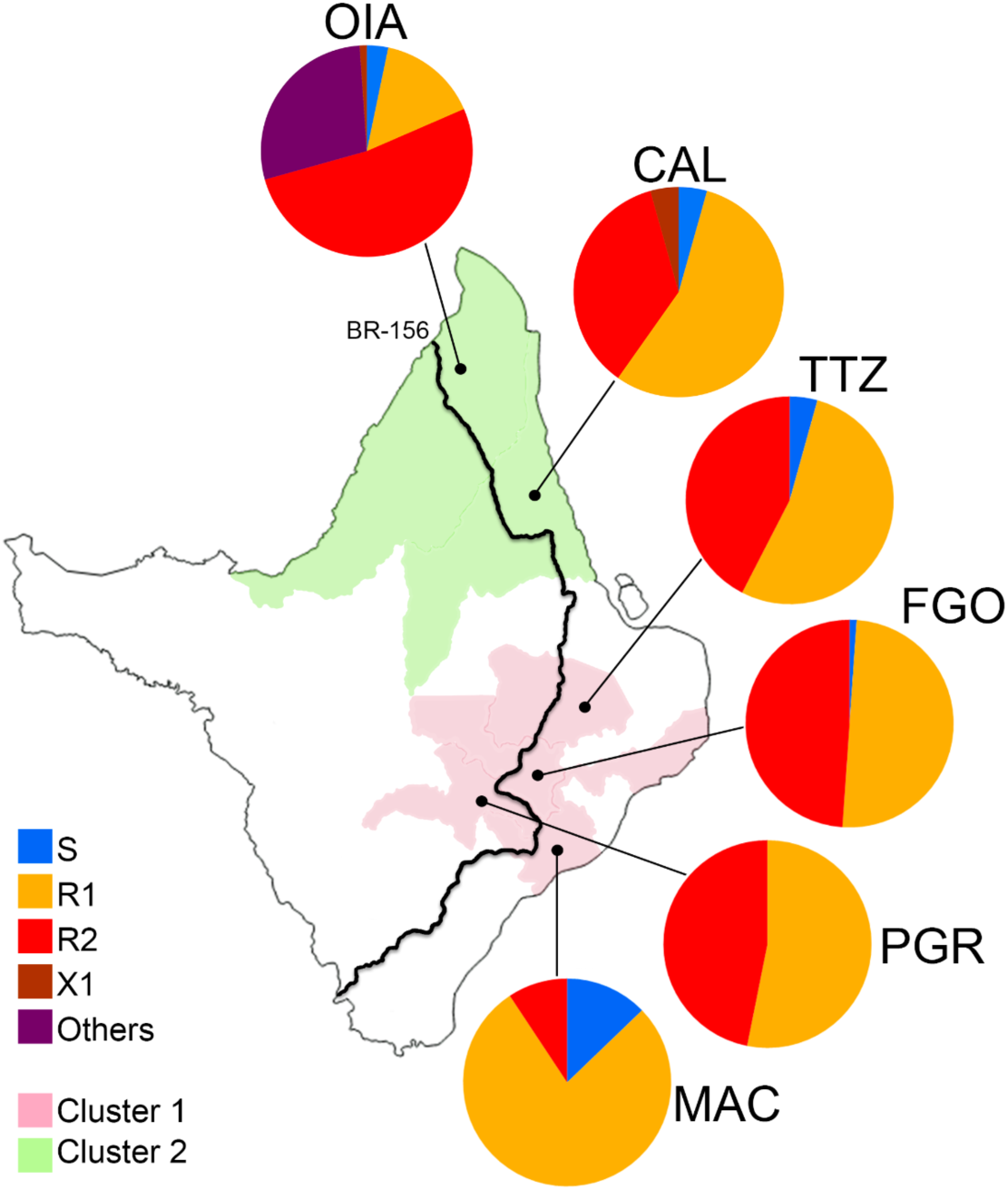
*Kdr* allelic frequencies and genetic clustering of *Aedes aegypti* populations from Amapá State, Brazil. *Kdr* allelic frequencies are indicated in pie charts for each respective locality. The localities are shaded according to their genetic clustering (K=2, see Fig 3). OIA: Oiapoque, CAL: Calçoene, TTZ: Tartarugalzinho, FGO: Ferreira Gomes, PGR: Porto Grande, MAC: Macapá. S: Wild-type *Na_v_S* (VVF), R1: *kdr Na_v_R1* (VVC), R2: *kdr Na_v_R2* (LIC), X1: *kdr Na_v_X1* (LVC), Others: alleles possibly composing the Y genotype: R2X1 (LVF/LIC) or X1X3 (LIF/LVC), see Fig 2B.

## Discussion

In this study, we investigated the frequency of *kdr* genotypes and the genetic structure of *Ae. aegypti* populations along a South-North transect of the Amapá State, starting from the capital city Macapá to Oiapoque, located on the border between Brazil and French Guiana. The Amapá State is particularly isolated from the other States and covered by >90% by the Amazon Rainforest. Our study showed that the dengue vector *Ae. aegypti* is divided into two well-defined genetic clusters in this State, and the *kdr* allelic composition roughly followed this clustering. We also found evidence of new *kdr* genotype arrangements in at least two of the six mosquito populations analyzed. Better understanding of the genetic structure of *Ae. aegypti* populations is relevant not only to better understand the possible dispersion of resistance genes in this region but also to guide decision-making for vector control and insecticide resistance management.

Historically in Brazil, the organophosphate malathion replaced pyrethroid insecticides around 2006-2009 [12]. However, malathion was never utilized in the Amapá State (operational constraints), and until recently, only pyrethroids were employed. The first cases of chikungunya in Brazil were recorded in 2014 in the cities of Oiapoque (Amapá State) and Feira de Santana (Bahia State), from two distinct events [25]. As a result, the use of pyrethroid insecticide (deltamethrin) was intensified in Oiapoque, which likely contributed to the increase in the selection pressure on resistance genes. Indeed, *Ae. aegypti* populations collected from Amapá State in 2014/2015 showed high resistance to deltamethrin with estimated RR_50_ of 46.4 and 143.9 in Macapá and Oiapoque, respectively [26]. Recently, Cielo has been used in ULV applications since 2019, and Spinosad was introduced in Amapá State in 2021 (*personal communication – Health Secretariat of Amapá State*). In French Guiana, deltamethrin is the only adulticide authorized in vector control campaigns since 2011 [48].

Here, we observed that all *Ae. aegypti* populations from Amapá presented high frequencies of *kdr* alleles, considering the three important SNPs in the voltage-gated sodium channel gene (*Na_v_*): V410L, V1016I and F1534C. Interestingly, we found two additional *kdr* genotypes composition or arrangement, LL+VI+CC in Calçoene and Oiapoque, and LL+VI+FC in Oiapoque. These distinct genotypes were probably present in previous surveys; however, they were not detected since the V410L had not been investigated in mosquitoes from Amapá [16]. The LL+VI+CC (R2X1) was previously observed in populations from Peru (Palomino, 2022 - unpublished data), Colombia [49], Mexico [50], and USA [51]. The LL+VI+FC (Y) was also previously observed in populations from the countries mentioned above, and in distinct Brazilian regions, such as Natal, Salvador, Vitória and Brasília [16]. Regardless the location, their frequencies were generally low: varying from 1.1% in the USA to 5.8% in Colombia (R2X1), and from 0.2% in Peru to 2.5% in the USA (Y). In Amapá the frequencies were 28.3% (Y) and 2.2% (R2X1) in Oiapoque, and 8.7% (R2X1) in Calçoene. Complementary studies conducted by our research team suggest that these genotypes may result from gene duplications, hence maintaining distinct copies of the *Na_v_* gene in the same chromosome [52,18,16]. Their relationship with pyrethroid resistance requires however further investigation.

We also showed that the *kdr Na_v_R2* allelic frequency remained very high in Oiapoque (67% in 2014/2015 [26], 68.5% in 2018 [16] and 52.2% in 2020) and it increased significantly in Calçoene (from 7.8% in 2018 [16] to 35.9% in 2020). While the *kdr Na_v_R2* was absent in Macapá in 2014/2015 [26], it has been found at low frequency (5.6%) in 2018 [16], and at 9.3% in 2020, as shown in this study. The homozygous R2R2 is still absent from Macapá but it has been reported in Calçoene for the first time. In the localities of Ferreira Gomes, Porto Grande and Tartarugalzinho *kdr Na_v_R2* varied from 42.4 to 48.9% and the R2R2 genotype ranged from 21.7 to 31.1%. It is worth noting that all localities along the BR-156 road had higher *kdr Na_v_R2* frequency than Macapá, which is the State capital and therefore highly connected to these localities. Considering the genetic cost of the *kdr Na_v_R2* under insecticide-free environment [18], we assume that the insecticide pressure by pyrethroids might be lower in Macapá compared to other cities, hence prevailing the introduction and establishment of the *kdr Na_v_R2* in *Ae. aegypti* in the State capital.

In addition, the decrease of the wild-type *Na_v_S* allele in all populations is of great concern. In Macapá, the frequency of the S allele was 15.7% in 2014/2015 [26], 17.8% in 2018 [16] and 12.8% in 2020. We also showed that the wild-type allele *Na_v_S* is now absent in Porto Grande and less than 13% in other populations tested. It is worth noting that the *Na_v_*S allele was not detected in any *Ae. aegypti* population from the Amazonian region in the previous nationwide surveillance (2018) [16].

Regarding the *kdr* alleles nucleotide composition, we showed that the sequences of R1R1 and R2R2 (homozygous) samples from Macapá and Oiapoque were similar to the ones previously described [17,53]. Evidence shows that at least two haplotypes emerged independently with the 1534C *kdr* mutation and one haplotype with the *kdr* mutation 1016I [17]. The 1534C is present in the herein called *kdr* alleles *Na_v_R1* (VVC) and *Na_v_R2* (LIC) and the 1016I in the *kdr Na_v_R2* in *Ae. aegypti* Brazilian populations [16].

Overall, we suspect that the high resistance to pyrethroids in Oiapoque is associated with a higher frequency of *kdr Na_v_R2* and the occurrence of rare *kdr* genotypes, as well as to the higher expression of detoxifying genes. Indeed, *Ae. aegypti* populations collected in the transborder city of Saint Georges de Oiapoque in French Guiana showed marked amplification of several *CYP6s* and *CYP9Js* playing a role in pyrethroid resistance [54]. We then cannot discard the possibility that these genes may be present in the Oiapoque population. Indeed Oiapoque (AP-Brazil) and Saint-Georges (French Guiana) are separated by a river which probably does not avoid the intense gene flow of *Ae. aegypti* between both sides, facilitated by the transportation of people and goods, which directly impacts the genetic structure of the mosquito [24]. Further work is needed to assess the detoxification pathway and genomic changes underlying the resistance mechanism of *Ae. aegypti* in the transborder area between Brazil and France that exhibits extremely high levels of resistance to all public health insecticides.

Finally, we analyzed the genetic structure and gene flow between *Ae. aegypti* from Amapá using 12 microsatellites markers. We showed that the Fst values among the six populations of *Ae. aegypti* were higher in the pairs with Oiapoque, which was isolated in a single genetic group in the Bayesian analysis (k=2) and clearly isolated in the DAPC. Calçoene, the closest city to Oiapoque, was also represented in an isolated group in the DAPC. The isolation by distance (IBD), confirmed by the Mantel test, and the dense Amazon Rainforest remain the most likely explanation for the genetic structure and *kdr* frequency differences among the populations alongside the BR-156 Road. Interestingly, Oiapoque presented the lowest Fst and the highest Nm with Macapá, that are yet distant from 580 Km. This can be explained by the fact that Macapá is the State capital, and the flow of people and goods between those cities is very intense, hence facilitating the passive transportation of *Ae. aegypti*. Other studies corroborated the relationship between gene flow in *Ae. Aegypti* and passive transportation through roads and waterways [55–57].

Based on the use of microsatellite markers, some studies concluded that the reinfestation of *Ae. aegypti* populations in Brazil after the eradication programme of the 1970s may have come from at least two distinct genetically differentiated groups, i.e. one from Venezuela to northern Brazil, and one from the Caribbean populations to the southeast of Brazil [58,59]. From this perspective, the Amapá State should have been invaded by *Ae. aegypti* from the two genetically distinct groups: in the north (Oaipoque and Calçoene) by the ‘Caribbean group’ through French Guiana, and in the south (Macapá, Ferreira Gomes, Porto Grande and Tartarugalzinho) by the ‘Venezuelan group’ through other Amazonian Brazilian locality, probably Pará State.

In this study, we used two types of molecular genotyping, *kdr* and microsatellite markers, to address distinct questions. The *kdr* SNPs were genotyped to obtain their genotypic frequencies in the populations. As *kdr* is under strong selection pressure, it is not an accurate marker to infer the population genetic structure, which was achieved by neutral microsatellite genotyping. By combining these two analyses, we observed that *kdr* alleles are spreading, likely by the *Ae. aegypti* ability to disperse itself and its eggs passively, however, without disrupting the overall genetic structure of the isolated populations. Further similar studies should be conducted over time, including eco-epidemiological data, to better understand the evolution of insecticide resistance of *Ae. aegypti* in the region.

## Conclusions

Microsatellites and *kdr* genotyping analyses evidenced genetic differences among *Ae. aegypti* populations relatively isolated by a dense forest, connected by an only principal road. We found that rare *kdr* genotypes are increasing in frequency in some regions of Amapá State and this relationship with pyrethroid resistance deserves further investigation. More additional work is also needed to better understand the environmental and landscape determinants involved in the evolution and spatial distribution of insecticide resistance including *kdr* mutations in the dengue vector *Aedes aegypti* in Amapá.

## Acknowledgment

We thank all staff from the IEPA (Instituto de Pesquisas Científicas e Tecnológicas do Estado do Amapá) for their assistance with mosquito collection. We thank the “Rede de Plataformas Tecnológicas Fiocruz (RPT/Fiocruz/RJ) – DNA Sequencing Platform” for their invaluable in this research and CNPq for investing in Barbara Souza’s graduation, making this study possible.

## Supporting information

**S1 Table. Primer and probe sequences for the SNPs V410L, V1016I and F1534C *kdr* in *Aedes aegypti*.**

**S2 Table. Primers used to amplify the IS6, IIS6 and IIIS6 fragments of the voltage-gated sodium channel gene (*Na_v_*).**

**S3 Table. Primers for microsatellite genotyping in *Aedes aegypti* populations. S4 Table. Frequencies of *kdr* genotypes in *Aedes aegypti* populations in the Amapá State, considering the *Na_v_* SNPs V410L, V1016I, F1534C.**

**S5 Table. Frequencies of *kdr* alleles in *Aedes aegypti* populations in the Amapá State, considering the *Na_v_* SNPs V410L, V1016I, F1534C.**

**S6 Table. Allele frequencies of the 12 microsatellite loci in *Aedes aegypti* populations in Amapá.**

**S7 Table. Analysis of the Genetic Diversity of *Aedes aegypti* using 12 microsatellite loci.**

**S8 Table. Analysis of linkage disequilibrium of *Aedes aegypti* populations in Amapá.**

**S9 Table. Population pairwise Fst value for *Aedes aegypti* population studied. S10 Table. Analysis of molecular variance (AMOVA) of Amapá populations.**

**S1 Fig. Frequency of *kdr* genotypes for each *Aedes aegypti* population in the Amapá State**

**S2 Fig. Evanno plot derived from STRUCTURE HARVESTER for detecting number of genetic clusters.**

## References

1. Girard M, Nelson CB, Picot V, Gubler DJ. Arboviruses: A global public health threat. Vaccine. 2020;38: 3989–3994. doi:10.1016/J.VACCINE.2020.04.011

2. PAHO. Pan American Health Organization. Epidemiological Update Dengue and other Arboviruses. 2020. Available: http://www.paho.org•©PAHO/WHO,2020

3. MS. Ministério da Saúde. Monitoramento dos casos de arboviroses até a semana epidemiológica 52 de 2022. 2023.

4. PAHO. Pan American Health Organization. Epidemiological Alert Yellow Fever Situation Summary. 2022. Available: https://bit.ly/2Gn9lzI

5. Nunes PCG, Daumas RP, Sánchez-Arcila JC, Nogueira RMR, Horta MAP, Dos Santos FB. 30 years of fatal dengue cases in Brazil: a review. BMC Public Health. 2019;19. doi:10.1186/S12889-019-6641-4

6. Achee NL, Gould F, Perkins TA, Reiner RC, Morrison AC, Ritchie SA, et al. A critical assessment of vector control for dengue prevention. PLoS Negl Trop Dis. 2015;9. doi:10.1371/JOURNAL.PNTD.0003655

7. Dusfour I, Vontas J, David J-P, Weetman D, Fonseca DM, Corbel V, et al. Management of insecticide resistance in the major Aedes vectors of arboviruses: Advances and challenges. PLoS Negl Trop Dis. 2019;13: e0007615. doi:10.1371/journal.pntd.0007615

8. WHO. World Health Organization. Global vector control response 2017–2030. 2017.

9. Moyes CL, Vontas J, Martins AJ, Ng LC, Koou SY, Dusfour I, et al. Contemporary status of insecticide resistance in the major Aedes vectors of arboviruses infecting humans. PLoS Negl Trop Dis. 2017;11: e0005625. doi:10.1371/journal.pntd.0005625

10. Corbel V, Achee NL, Chandre F, Coulibaly MB, Dusfour I, Fonseca DM, et al. Tracking Insecticide Resistance in Mosquito Vectors of Arboviruses: The Worldwide Insecticide resistance Network (WIN). PLoS Negl Trop Dis. 2016;10. doi:10.1371/JOURNAL.PNTD.0005054

11. Moyes CL, Vontas J, Martins AJ, Ng LC, Koou SY, Dusfour I, et al. Correction to: Contemporary status of insecticide resistance in the major aedes vectors of arboviruses infecting humans (PLoS Negl Trop Dis). PLoS Negl Trop Dis. 2021;15: 1–2. doi:10.1371/journal.pntd.0009084

12. Valle D, Bellinato DF, Viana-Medeiros PF, Lima JBP, Martins Junior ADJ. Resistance to temephos and deltamethrin in Aedes aegypti from Brazil between 1985 and 2017. Mem Inst Oswaldo Cruz. 2019;114. doi:10.1590/0074-02760180544

13. Campos KB, Martins AJ, Rodovalho C de M, Bellinato DF, Dias L dos S, Macoris M de L da G, et al. Assessment of the susceptibility status of Aedes aegypti (Diptera: Culicidae) populations to pyriproxyfen and malathion in a nation-wide monitoring of insecticide resistance performed in Brazil from 2017 to 2018. Parasit Vectors. 2020;13. doi:10.1186/S13071-020-04406-6

14. Williamson MS, Martinez-Torres D, Hick CA, Devonshire AL. Identification of mutations in the houseflypara-type sodium channel gene associated with knockdown resistance (*kdr*) to pyrethroid insecticides. Mol Gen Genet. 1996;252: 51–60. doi:10.1007/BF02173204

15. Haddi K, Tomé HVV, Du Y, Valbon WR, Nomura Y, Martins GF, et al. Detection of a new pyrethroid resistance mutation (V410L) in the sodium channel of Aedes aegypti: a potential challenge for mosquito control. Scientific Reports 2017 7:1. 2017;7: 1–9. doi:10.1038/srep46549

16. Melo Costa M, Campos KB, Brito LP, Roux E, Melo Rodovalho C, Bellinato DF, et al. *Kdr* genotyping in Aedes aegypti from Brazil on a nation-wide scale from 2017 to 2018. Sci Rep. 2020;10. doi:10.1038/S41598-020-70029-7

17. Cosme LV, Gloria-Soria A, Caccone A, Powell JR, Martins AJ. Evolution of *kdr* haplotypes in worldwide populations of Aedes aegypti: Independent origins of the F1534C *kdr* mutation. PLoS Negl Trop Dis. 2020;14: e0008219. doi:10.1371/journal.pntd.0008219

18. Brito LP, Carrara L, Freitas RMD, Lima JBP, Martins AJ. Levels of Resistance to Pyrethroid among Distinct *kdr* Alleles in Aedes aegypti Laboratory Lines and Frequency of *kdr* Alleles in 27 Natural Populations from Rio de Janeiro, Brazil. Biomed Res Int. 2018;2018. doi:10.1155/2018/2410819

19. Kushwah RBS, Dykes CL, Kapoor N, Adak T, Singh OP. Pyrethroid-resistance and presence of two knockdown resistance (kdr) mutations, F1534C and a novel mutation T1520I, in Indian Aedes aegypti. PLoS Negl Trop Dis. 2015;9. doi:10.1371/JOURNAL.PNTD.0003332

20. Ur Rahman R, Souza B, Uddin I, Carrara L, Paulo Brito L, Melo Costa M, et al. Insecticide resistance and underlying targets-site and metabolic mechanisms in Aedes aegypti and Aedes albopictus from Lahore, Pakistan. Scientific Reports |. 123AD;11: 4555. doi:10.1038/s41598-021-83465-w

21. Donnelly MJ, Corbel V, Weetman D, Wilding CS, Williamson MS, Black WC 4th. Does *kdr* genotype predict insecticide-resistance phenotype in mosquitoes? Trends Parasitol. 2009;25: 213–219. doi:10.1016/j.pt.2009.02.007

22. Roiz D, Wilson AL, Scott TW, Fonseca DM, Jourdain F, Müller P, et al. Integrated Aedes management for the control of Aedes-borne diseases. PLoS Negl Trop Dis. 2018;12: e0006845. doi:10.1371/journal.pntd.0006845

23. Suesdek L. Microevolution of medically important mosquitoes – A review. Acta Trop. 2019;191: 162–171. doi:10.1016/J.ACTATROPICA.2018.12.013

24. Salgueiro P, Restrepo-Zabaleta J, Costa M, Galardo AKR, Pinto J, Gaborit P, et al. Liaisons dangereuses: cross-border gene flow and dispersal of insecticide resistance-associated genes in the mosquito Aedes aegypti from Brazil and French Guiana. Mem Inst Oswaldo Cruz. 2019;114. doi:10.1590/0074-02760190120

25. Nunes MRT, Faria NR, de Vasconcelos JM, Golding N, Kraemer MUG, de Oliveira LF, et al. Emergence and potential for spread of Chikungunya virus in Brazil. BMC Med. 2015;13. doi:10.1186/s12916-015-0348-x

26. Costa MM. Avaliação da resistência a inseticidas e mecanismos selecionados em populações de Aedes aegypti Linnaeus 1762 (Diptera, Culicidae) da fronteira entre Brasil e Guiana Francesa. FUNDAÇÃO OSWALDO CRUZ/ INSTITUTO OSWALDO CRUZ. 2017.

27. Cosme LV, Lima JBP, Powell JR, Martins AJ. Genome-wide Association Study Reveals New Loci Associated With Pyrethroid Resistance in Aedes aegypti. Front Genet. 2022;13. doi:10.3389/FGENE.2022.867231/FULL

28. Martins J, Solomon SE, Mikheyev AS, Mueller UG, Ortiz A, Bacci M. Nuclear mitochondrial-like sequences in ants: Evidence from Atta cephalotes (Formicidae: Attini). Insect Mol Biol. 2007;16: 777–784. doi:10.1111/j.1365-2583.2007.00771.x

29. Kearse M, Moir R, Wilson A, Stones-Havas S, Cheung M, Sturrock S, et al. Geneious Basic: An integrated and extendable desktop software platform for the organization and analysis of sequence data. Bioinformatics. 2012;28: 1647–1649. doi:10.1093/bioinformatics/bts199

30. Brown JE, Mcbride CS, Johnson P, Ritchie S, Paupy C, Bossin H, et al. Worldwide patterns of genetic differentiation imply multiple “domestications” of Aedes aegypti, a major vector of human diseases. Proceedings of the Royal Society B: Biological Sciences. 2011;278: 2446–2454. doi:10.1098/rspb.2010.2469

31. Slotman MA, Kelly NB, Harrington LC, Kitthawee S, Jones JW, Scott TW, et al. Polymorphic microsatellite markers for studies of Aedes aegypti (Diptera: Culicidae), the vector of dengue and yellow fever. Mol Ecol Notes. 2007;7: 168–171. doi:10.1111/j.1471-8286.2006.01533.x

32. Schuelke M. An economic method for the fluorescent labeling of PCR fragments A poor man ’ s approach to genotyping for research and high-throughput diagnostics. Prism. 2000;18: 1–2. Available: https://www.ncbi.nlm.nih.gov/pubmed/10657137

33. Rousset F, Raymond M. GENEPOP (version 3.3): population genetics software for exact tests and ecumenicism. J Hered. 1995;86: 248–249.

34. Rousset F. genepop’007: a complete re-implementation of the genepop software for Windows and Linux. Mol Ecol Resour. 2008;8: 103–106. doi:10.1111/J.1471-8286.2007.01931.X

35. Van Oosterhout C, Hutchinson WF, Wills DPM, Shipley P. micro-checker: software for identifying and correcting genotyping errors in microsatellite data. Mol Ecol Notes. 2004;4: 535–538. doi:10.1111/J.1471-8286.2004.00684.X

36. Peakall R, Smouse PE. GenALEx 6.5: Genetic analysis in Excel. Population genetic software for teaching and research-an update. Bioinformatics. 2012;28: 2537–2539. doi:10.1093/bioinformatics/bts460

37. Kalinowski ST. hp-rare 1.0: a computer program for performing rarefaction on measures of allelic richness. Mol Ecol Notes. 2005;5: 187–189. doi:10.1111/J.1471-8286.2004.00845.X

38. Excoffier L, Lischer HEL. Arlequin suite ver 3.5: a new series of programs to perform population genetics analyses under Linux and Windows. Mol Ecol Resour. 2010;10: 564–567. doi:10.1111/J.1755-0998.2010.02847.X

39. Chapuis M-P, Estoup A. Microsatellite null alleles and estimation of population differentiation. Mol Biol Evol. 2007;24: 621–631. doi:10.1093/molbev/msl191

40. Pritchard JK, Stephens M, Donnelly P. Inference of Population Structure Using Multilocus Genotype Data. Genetics. 2000;204: 391–393. doi:10.1534/genetics.116.195164

41. Earl DA, vonHoldt BM. STRUCTURE HARVESTER: A website and program for visualizing STRUCTURE output and implementing the Evanno method. Conserv Genet Resour. 2012;4: 359–361. doi:10.1007/s12686-011-9548-7

42. Evanno G, Regnaut S, Goudet J. Detecting the number of clusters of individuals using the software STRUCTURE: A simulation study. Mol Ecol. 2005;14: 2611–2620. doi:10.1111/j.1365-294X.2005.02553.x

43. Jakobsson M, Rosenberg NA. CLUMPP: a cluster matching and permutation program for dealing with label switching and multimodality in analysis of population structure. Bioinformatics. 2007;23: 1801–1806. doi:10.1093/bioinformatics/btm233

44. Rosenberg NA. distruct: a program for the graphical display of population structure. Mol Ecol Notes. 2004;4: 137–138. doi:10.1046/J.1471-8286.2003.00566.X

45. Jombart T, Bateman A. adegenet: a R package for the multivariate analysis of genetic markers. Bioinformatics. 2008;24: 1403–1405. doi:10.1093/BIOINFORMATICS/BTN129

46. Jombart T, Devillard S, Balloux F. Discriminant analysis of principal components: A new method for the analysis of genetically structured populations. BMC Genet. 2010;11: 1–15. doi:10.1186/1471-2156-11-94/FIGURES/9

47. R Core Team (2018). A Language and Environment for Statistical Computing. R Foundation for Statistical Computing, Vienna. 2018. Available: https://www.r-project.org/

48. Epelboin Y, Chaney SC, Guidez A, Habchi-Hanriot N, Talaga S, Wang L, et al. Successes and failures of sixty years of vector control in French Guiana: what is the next step? Mem Inst Oswaldo Cruz. 2018;113: 1–10. doi:10.1590/0074-02760170398

49. Granada Y, Mejía-Jaramillo AM, Zuluaga S, Triana-Chávez O. Molecular surveillance of resistance to pyrethroids insecticides in Colombian Aedes aegypti populations. PLoS Negl Trop Dis. 2021;15. doi:10.1371/JOURNAL.PNTD.0010001

50. Saavedra-Rodriguez K, Maloof FV, Campbell CL, Garcia-Rejon J, Lenhart A, Penilla P, et al. Parallel evolution of vgsc mutations at domains IS6, IIS6 and IIIS6 in pyrethroid resistant Aedes aegypti from Mexico. Scientific Reports 2018 8:1. 2018;8: 1–9. doi:10.1038/s41598-018-25222-0

51. Hernandez JR, Liu S, Fredregill CL, Pietrantonio PV. Impact of the V410L *kdr* mutation and co-occurring genotypes at *kdr* sites 1016 and 1534 in the VGSC on the probability of survival of the mosquito Aedes aegypti (L.) to Permanone in Harris County, TX, USA. PLoS Negl Trop Dis. 2023;17: e0011033. doi:10.1371/JOURNAL.PNTD.0011033

52. Martins AJ, Brito LP, Linss JGB, Rivas GB da S, Machado R, Bruno RV, et al. Evidence for gene duplication in the voltage-gated sodium channel gene of Aedes aegypti. Evolution, medicine, and public health. 2013;2013: 148–160. doi:10.1093/EMPH/EOT012

53. Itokawa K, Furutani S, Takaoka A, Maekawa Y, Sawabe K, Komagata O, et al. A first, naturally occurring substitution at the second pyrethroid receptor of voltage-gated sodium channel of Aedes aegypti. Pest Manag Sci. 2021;77: 2887–2893. doi:10.1002/PS.6324

54. Faucon F, Gaude T, Dusfour I, Navratil V, Corbel V, Juntarajumnong W, et al. In the hunt for genomic markers of metabolic resistance to pyrethroids in the mosquito Aedes aegypti: An integrated next-generation sequencing approach. PLoS Negl Trop Dis. 2017;11: e0005526. doi:10.1371/JOURNAL.PNTD.0005526

55. Sá ELR de, Rodovalho C de M, Sousa NPR de, Sá ILR de, Bellinato DF, Dias L dos S, et al. Evaluation of insecticide resistance in *Aedes aegypti* populations connected by roads and rivers: the case of Tocantins state in Brazil. Mem Inst Oswaldo Cruz [Internet]. 2019;114:e180318. doi: 10.1590/0074-02760180318

56. Díaz-Nieto LM, Chiappero MB, Díaz de Astarloa C, Maciá A, Gardenal CN, Berón CM. Genetic Evidence of Expansion by Passive Transport of Aedes (Stegomyia) aegypti in Eastern Argentina. PLoS Negl Trop Dis. 2016;10. doi:10.1371/JOURNAL.PNTD.0004839

57. Guagliardo SAJ, Lee Y, Pierce AA, Wong J, Chu YY, Morrison AC, et al. The genetic structure of Aedes aegypti populations is driven by boat traffic in the Peruvian Amazon. PLoS Negl Trop Dis. 2019;13. doi:10.1371/JOURNAL.PNTD.0007552

58. Monteiro FA, Shama R, Martins AJ, Gloria-Soria A, Brown JE, Powell JR. Genetic Diversity of Brazilian Aedes aegypti: Patterns following an Eradication Program. PLoS Negl Trop Dis. 2014;8. doi:10.1371/journal.pntd.0003167

59. Kotsakiozi P, Gloria-Soria A, Caccone A, Evans B, Schama R, Martins AJ, et al. Tracking the return of Aedes aegypti to Brazil, the major vector of the dengue, chikungunya and Zika viruses. PLoS Negl Trop Dis. 2017;11: e0005653. doi:10.1371/JOURNAL.PNTD.0005653

